# Slightly different metabolomic profiles are associated with high or low weights duck foie gras

**DOI:** 10.1101/2021.08.04.455120

**Authors:** Bara Lo, Nathalie Marty-Gasset, Hélène Manse, Cécile Canlet, Renaud Domitile, Hervé Rémignon

## Abstract

Understanding the evolution of fatty liver metabolism of ducks is a recurrent issue for researchers and industry. Indeed, the increase in weight during the overfeeding period leads to an important change in the liver metabolism. However, liver weight is highly variable at the end of overfeeding within a batch of animals reared, force-fed and slaughtered in the same way. For this study, we performed a proton nuclear magnetic resonance (^1^H-NMR) analysis on two classes of fatty liver samples, called low-weight liver (weight between 550 and 599 g) and high-weight liver (weight above 700 g). The aim of this study was to identify the differences in metabolism between two classes of liver weight (low and high). Firstly, the results showed that increased liver weight is associated with higher glucose uptake leading to greater lipid synthesis. Secondly, this increase is probably also due to a decline in the level of export of triglycerides and cholesterol from the liver by maintaining them at high hepatic concentration levels. Finally, the increase in liver weight could lead to a significant decrease in the efficiency of aerobic energy metabolism associated with a significant increase in the level of oxidative stress.

## Introduction

France generates about 80% of the world production of “Foie gras”. This is mainly composed of mule duck “foie gras”, which are interspecific hybrids produced by crossing a common female duck (*Anas Platyrhynchos*) with a Muscovy male duck (*Caïrina Moschata*). To obtain this product, male mule ducks are overfed for up to 12 days after a rearing period of about 12 weeks. During the overfeeding period, the ducks are fed ground corn mixed with water to form a flour that is more digestible. This diet is hypercaloric because corn is composed of 64% starch [1] and finally leads to a hepatic steatosis. Indeed, glucose absorbed by the hepatocytes is transformed into glycogen and when it reaches about 5% of the hepatic mass, the surplus is redirected to pathways leading to the synthesis of fatty acids which will be esterified into triglycerides to be exported to the adipose tissue in the form of very low-density lipoproteins [2,3]. Moreover, when the storage capacity of peripheral tissues reaches saturation, the reabsorption of circulating lipids allows the liver to also become the main fat storage site during this process [4]. Thus, at the end of the overfeeding period, a massive storage of lipids in the hepatocytes is observed and largely contributes to a huge increase in the weight of the liver (x10 between the beginning and the end of the overfeeding period). Livers issued from overfed ducks must have a weight greater than 300 g to be legally called « foie gras » [5] but their weights can reach more than 700 g at the end of this period [6]. To achieve this high level of synthesis and storage of lipids, several changes in various metabolic activities of the liver have also been observed such as various levels of hypoxia and of oxidative stress, modifications of cellular proteins metabolisms, cytoskeleton and extracellular matrix reorganizations and even cellular deaths by apoptosis [7–9]. However, a large intra-batch variability of final liver weights is generally observed by professionals despite similar intake amounts throughout the overfeeding process. Recent studies on the proteins fraction have been carried out on two classes of liver weights, light (weight between 550 and 600 g) and heavy (weight over 700 g) and showed marked differences in energy metabolism, oxidative stress, lipids and proteins metabolisms and in the extracellular membrane reorganization [9].

To get a complementary overview of the involvement of main hepatic metabolites in this process, we performed a new comparison with a metabolomic approach using 1H-NMR between ducks of the same breed, raised and overfed under the same conditions but presenting different liver weights at the end of the overfeeding period. The aim of this study is therefore to identify which essential metabolites significantly vary between low and high weight livers and how they inform us about differences in metabolism activities between those two groups of weight livers.

## Material and methods

A flock of about 2000 male mule ducks (*Caïrina moschata x Anas platyrhynchos*) was reared for 12 weeks according to standard commercial rules. Then, birds were overfed for 20 meals (twice a day during 10 days) according to a standard overfeeding program based on moistened corn flour. At the beginning of the overfeeding period, ducks received a quantity of 225 g/meal which was progressively increased to a final value of 510 g for the last meal. As a whole, during the overfeeding period, ducks ingested an average quantity of 8,8 kg of feed. Ducks were slaughtered approximately 11h hours after the last meal in a commercial slaughterhouse. At 20 min *post-mortem,* 40 livers were randomly sampled according to their weights to create 2 experimental groups: 20 livers weighing between 550 g and 600 g (low weight livers or LWL groups) and 20 livers with a weight over than 700 g (high weight livers or HWL group). From all those livers, 50 g were collected from the median lobe and directly frozen in liquid nitrogen before storage at - 80 °C. The rest of the liver was cooled to 4°C in cooled air. Then, Near Infra-Red Spectra (NIRS) were performed in absorbance from 250 to 2500 nm with an interval of 1 nm with the Labspec® 5000 Pro spectrometer (ASD Inc., Boulder (CO), USA), on six independent points of the surface of each liver, to predict the raw biochemical characteristics of the samples.

### Biochemical analyses

For all samples, the raw biochemical (i.e. dry matter, total lipids and total nitrogen) contents were determined from NIRS spectra according to the method described by Marie-Etancelin et al. (2014) [10]. 16 samples (8/groups) were then selected for being representatives of the whole 40 samples and further analyzed for metabolomics. The total amount of proteins was determined by the formula: (% Proteins = % total nitrogen × 6.25).

### Metabolomic analysis

#### Metabolite’s extraction and 1H-NMR Analysis

Polar metabolites were extracted from samples according to a modified protocol from Beckonert et al., (2007) [11]. Briefly, 0.25 g of crushed liver was added to methanol in a ratio 1:4 (w/v) and water in a ratio 1:0.85 (w/v) at 4°C and homogenized in Fastprep®-24 Instrument (MP Biomedicals, USA). Subsequently, dichloromethane in a ratio 1:2 (w/v) was added twice and mixed between each addition. Then, water was added in a ratio 1:2 (w/v). After 15 min at 4°C, the mixture was centrifuged at 1 000g at 4°C for 15 min. The upper phase, composed of hydrophilic metabolites, was collected in new polypropylene tubes and evaporated with a vacuum concentrator (Concentrator Plus, Eppendorf, Hamburg, Germany). Then, each sample was diluted into 650 μl of NMR buffer phosphate pH=7 in deuterated water (D_2_O) with sodium trimethylsilylpropionate (17,2g TMSP for 100ml), and stored at −20°C until NMR (Nuclear Magnetic Resonance) analysis. Thereafter, the tube was thawed homogenized and centrifuged for 15 minutes at 5 350 g. Finally, 600 μl were transferred in a NMR tube.

The ^1^H-NMR analysis of polar metabolites was done at MetaToul-AXIOM platform (http://www.metatoul.fr/) equipped with a Bruker Avance III HD NMR spectrometer at a proton resonance frequency of 600 Mhz. Topspin (V2.1, Bruker, Biospin, Munich, Germany) was used and the NOESYPR1D spin echo pulse sequence was applied to attenuate the signal in the water [12]. The spectra were acquired at 300 K and each spectrum collected with 32 K data points, 16 dummy scans and 512 scans. After Fourier’s transformation, all spectra were manually phased, baseline corrected, and chemicals shifts were calibrated to TMSP at 0 ppm. To confirm the chemical structure of metabolites of interest, 2D ^1^H - ^1^H – COSY (COrrelation SpectroscopY) and 2D ^1^H - ^13^C HSQC (Heteronuclear Single Quantum Coherence spectroscopy) NMR experiment was performed on one sample.

#### Spectra pre-processing and statistical analysis

In a nutshell, the metabolite method involved the use of the ASICS R software package (R software package version 4.0.2 https://bioconductor.org/packages/ASICS/) to convert ^1^H-NMR spectra into a metabolite relative concentration table. It contains an automatic identification method and quantifies the metabolites in the complex ^1^H-NMR spectrum based on its unique peak shape [13,14]. The metabolite database used consists of the spectra of 190 pure metabolites described by Lefort et al., (2019), therefore only metabolites included in this list will be identified and quantified [15]. The zones corresponding to water (5.0 - 4.7 ppm), dichloromethane (5.5 - 5.44 ppm) and methanol (3.38 - 3.34 ppm) were excluded. Finally, a table of metabolite relative concentration in column and sample number in rows was generated.

Statistical analyses were performed with the R software (version 4.0.2). First, a partial least squares discriminant analysis (PLS-DA) was performed with MixOmics R package (R package version 4.0.2. https://CRAN.R-project.org/package=mixOmics) as described by Lê Cao et al., (2011) and the ropls R package (R package version 4.0.2. https://www.bioconductor.org/packages/release/bioc/html/ropls.html) was used to highlight the metabolites useful for distinguishing the two (LWL and HWL) originals groups of livers [16,17]. The quality of fit of the models was estimated by the proportion of cumulative explained variance (R^2^) for the variables X (X = metabolites) and the variable Y (Y = Liver weight) and by the predictive ability of the model (Q^2^). First of all, Q^2^ is calculated by a cross-validation, and predictive data are then compared with the original ones and the sum of squared errors calculated for the whole dataset. Then, the Root Mean Square Error of Estimation (RMSEE) was computed and indicated the fit of the observations to the model. When its value is zero, it means that the correlation coefficient is 1 and implies that all points lie on the regression line. So, the lower its value, the better the quality of the model [18]. A latent variable is only included in the PLS model if its Q^2^ value is greater than or equal to 0.0975 [19]. Finally, the model parameters of the original data (R2Y and Q2) and those of all models for 500 datasets when Y (liver weight) is randomly permuted with an unchanged X (metabolite) matrix (pR^2^Y and pQ^2^) were compared. The model was validated only if all parameter values of the permuted model were much lower than those of the original model and the intercept of the regression line passing through the scatterplot was less than 0. For the final model, only variables with Variable Importance in Projection (VIP) values greater than 1 were considered to be discriminatory. Then, the Student - Wilcoxon’s mean comparison analysis was performed and only metabolites with a p-value <0.05 were considered as significantly different with multiple testing correction false-discovery rate (FDR).

The experiments described here fully comply with legislation on research involving animal subjects according to the European Communities Council directive of November 24, 1986 (86/609/EEC). Investigators were certificated by the French governmental authority (agreement no. R-31-ENVT-F1-12) and by the Federation of European Laboratory Animal Science Associations (agreement no. F011/05) for carrying out those experiments.

## Results

### Biochemicals analysis

Liver weights and total protein contents from the two experimental groups were largely different (p-value < 0.001, Table 1). On the contrary, no significant differences (p-value > 0, 05) were observed for dry matter and total lipids contents between the two studied groups. These results are in accordance with observations made in previous studies [6,20].

**Table 1.** Liver weights (LW), gross chemical composition of livers from the two studied groups (n = 8 samples /groups).

|  | LWL | HWL | p. value |
| --- | --- | --- | --- |
| LW (g) | $581 \pm 18$ | $745 \pm 24$ | *** |
| Lipids (% of dry matter) | $61.4 \pm 0.5$ | $61.6 \pm 0.7$ | NS |
| Dry Matter (% of raw weight) | $71.3 \pm 1.8$ | $71.4 \pm 1.5$ | NS |
| Proteins (% of dry matter) | $6.64 \pm 0.41$ | $5.76 \pm 0.38$ | *** |

Values are means ± SD. Within a line, p values indicate the level of significance (\* = $P \leq 0.05$ , \*\* = $P \leq 0.01$ , \*\*\* = $P \leq 0.001$ and NS = non significative).

### Metabolomic analysis

As a result, ASICS R software was used to identify a table of 30 metabolites with their intensities from the 96 spectra composed mainly of sugars and amino acids and their derivatives. With the PLS-DA, using one latent variable model, the cumulative parameters of the model were R^2^X = 0.21 and R^2^Y = 0.792 and the prediction of the model was Q^2^ = 0.587 (Table 2). Adding a second latent variable only allowed to gain 0,01 for Q^2^, so less than the expected value of 0.0975. Consequently, the model with one latent variable was kept as the final model [21]. The model values for R^2^Y(cum) and Q^2^(cum) were higher than those obtained after permutation (pR^2^Y(cum) = 0.002 and pQ^2^(cum) = 0.004) (Table 2). The final RMSEE value was considered as being low (= 0.244).

**Table 2.** Standard scaling of predictors and response.

| Component | $R^2X$ | $R^2X(cum)$ | $R^2Y$ | $R^2Y(cum)$ | $Q^2$ | $Q^2(cum)$ | RMSEE | $pR^2Y$ | $pQ^2$ |
| --- | --- | --- | --- | --- | --- | --- | --- | --- | --- |
| 1 | 0.21 | 0.21 | 0.792 | 0.792 | 0.587 | 0.587 | 0.244 | 0.002 | 0.004 |
| 2 | 0.19 | 0.4 | 0.074 | 0.866 | 0.01 | 0.597 | 0.203 | 0.022 | 0.002 |

Thanks to the PLS-DA, the distinction of the two groups of livers, based on their respective metabolomics profiles, was effective along the horizontal axis which corresponds to the first latent variable (Fig 1).

**Fig 1.**
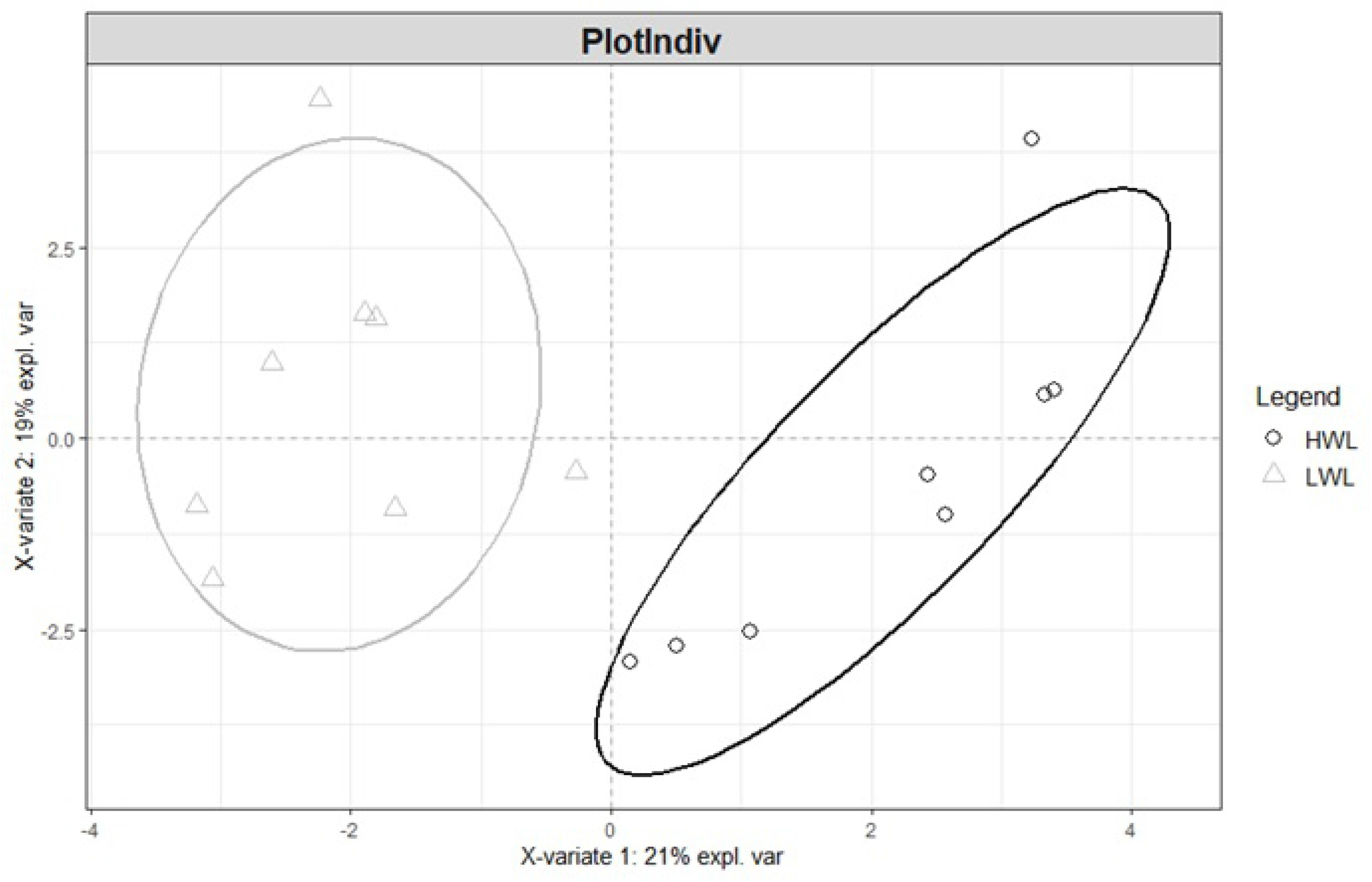
PLS-DA score plots according to the two first principal components for the two liver weight groups and their 30 variant metabolites (n = 8 livers /group).

Fourteen metabolites presented a VIP >1 and then represented about 50% of the initial 30 quantified metabolites (Table 3). Among this shortened list, 6 metabolites were identified as being the significant ones because they presented an FDR-corrected p-value < 0.05 (Table 3) corresponding to 20% of the initial quantified metabolites. Finally, calculations of the ratios between LWL and HWL for each significant metabolite indicate that 3 of them (creatine, lactate and L-glutathione.oxidized) were in higher amounts in the HWL than in the LWL group. On the contrary, 3 metabolites (choline chloride, D-glucose and ethanolamine) were more present in samples from the LWL group than in those from the HWL group (Table 3).

**Table 3.** List of the 30 metabolites identified in the samples from the LWL and/or the HLW groups. VIP values > 1 and FDR-corrected p-value < 0,05 allowed to identify the six significant LWL / HWL ratios which were reported in bold (n = 8 samples /groups).

| Metabolites | LWL/HWL | VIP | P.value |
| --- | --- | --- | --- |
| D-Sorbitol | 0.57 | 1.26 | 1.15E-01 |
| D-Maltose | 0.68 | 1.19 | 1.41E-01 |
| **Creatine** | **0.75** | **1.45** | **2.10E-02** |
| L-Glutamine | 0.76 | 1.07 | 1.06E-01 |
| **Lactate** | **0.81** | **1.87** | **1.11E-03** |
| **L-Glutathione-oxidized** | **0.83** | **1.66** | **5.96E-03** |
| 2-Oxoglutarate | 0.86 | 0.68 | 9.58E-01 |
| L-GlutamicAcid | 0.89 | 0.65 | 3.92E-01 |
| L-Alanine | 0.89 | 1.10 | 1.25E-01 |
| L-Serine | 0.89 | 0.86 | 2.70E-01 |
| Phosphocholine | 0.92 | 0.35 | 6.89E-01 |
| Myo-Inositol | 0.94 | 0.43 | 5.39E-01 |
| L-Glycine | 0.97 | 0.42 | 7.16E-01 |
| ArgininosuccinicAcid | 0.97 | 0.41 | 7.21E-01 |
| Taurine | 1.02 | 0.14 | 9.04E-01 |
| L-Aspartate | 1.03 | 0.24 | 8.36E-01 |
| L-Valine | 1.06 | 0.59 | 6.06E-01 |
| L-Carnitine | 1.06 | 0.64 | 3.99E-01 |
| L-Leucine | 1.07 | 0.70 | 4.46E-01 |
| L-Proline | 1.07 | 0.64 | 4.76E-01 |
| Betaine | 1.09 | 0.18 | 8.14E-01 |
| Succinate | 1.10 | 0.72 | 3.58E-01 |
| Uridine | 1.14 | 1.18 | 6.92E-02 |
| **Ethanolamine** | **1.16** | **1.44** | **2.73E-02** |
| AceticAcid | 1.18 | 1.13 | 8.98E-02 |
| Glycerophosphocholine | 1.18 | 1.11 | 9.08E-02 |
| Glycerol | 1.20 | 0.57 | 7.42E-02 |
| **D-Glucose** | **1.26** | **1.91** | **7.72E-04** |
| MalicAcid | 1.56 | 1.04 | 2.94E-01 |
| **CholineChloride** | **2.98** | **1.55** | **1.22E-02** |

## Discussion

During the overfeeding period, the high-caloric diet provided to the animals leads to a large amount of lipids synthesis in the liver and a large part of these lipids remains stored in macrovacuoles located in the cytoplasm of hepatocytes [22]. This process, physiologically defined as liver steatosis, results in an increase in the liver weight from about 80 g to over 700 g [6,8]. In parallel, the percentage of lipids stored in hepatocytes rose from 5% to more than 50% of the total dry matter (DM) [4,6].

The glucose, issued from the digestion of the starch which is present in large amounts in the corn, is at the origin of the huge increase in the synthesis of lipids because it is the main precursor of triacylglycerols synthesis in the liver of ducks [23]. Indeed, according to Evans (1972), in birds, about 75% of the glucose entering into the hepatocytes is converted into lipids in the form of fatty acids [24]. Although starch intake increased with increasing amounts of feed at each meal, Mozduri et al., (2021) showed that the plasmatic glucose content was negatively correlated to liver weight (r = - 0.94) in force-fed ducks [25]. Therefore, a decrease in the amount of glucose present in the liver may indicate an increase of its utilization for an insitu synthesis of lipids and consequently the increase in the content of lipids in hepatocytes. Therefore, in the present experiment, the higher amount of glucose recorded in LWL than in HWL induces that lipids synthesis might be more active in HWL than in LWL.

Choline chloride is a salt of choline and is known to be an essential nutrient for poultry [26]. In fact, choline plays important roles in the synthesis of the phospholipids of the membranes and in the methyl-group metabolism [27]. According to Wen et al., (2014), as choline increases in the diet, weight gain and food consumption rose linearly or quadratically. However, maize is a choline poor diet [1], therefore a variation in the amount of choline between LWL and HWL livers might not be related to the feeding but to an increase in its biosynthesis in the liver. Several studies have shown that choline prevented excessive lipids accumulation and the incidence of fatty liver in animals [28–30]. This is largely due to the fact that, phosphatidylcholine, a metabolite derived from the choline, is required for the packaging and the export of triglycerides into very low density lipoproteins (VLDL) [31]. Then, if choline is present at a low level in the liver, less VLDL can be synthetized and consequently less lipids will be exported. This favors the development of the hepatic steatosis. In the present study, a greater amount of choline was reported in LWL samples. This probably indicates that LWL, at the end of the overfeeding period, were still able to synthetize large amounts of VLDL and thus to export a part of the lipids neo-synthetized by the hepatocytes. In consequence, the weight of LWL is lower than HWL but this also signifies that hepatocytes from LWL are probably in a more active metabolism than those from HWL which are finally forced to store lipids because of their incapacity to export them.

Ethanolamine is known to be a precursor of the phosphatidylethanolamine (PE) which is an important component of the cellular membranes. Shimada et al., (2003) compared rats fed an ethanolamine-enriched diet with control rats and observed that the supplementation with ethanolamine induced, in the liver, a significant reduction in cholesterol and triglyceride amounts and an increase in phospholipids [32]. According to Leonardi et al. (2009) and Pavlovic and Bakovic (2013) the reduction of phosphatidylethanolamine synthesis in the liver triggered an increase in fatty acid synthesis to be incorporated in triglycerides and thus favored the development of a hepatic steatosis [33,34]. In the present study, the amount of ethanolamine was higher in LWL than in HWL samples [35]. This indicates that HWL had less capacities to elaborate PE and then are more oriented in the synthesis of triglycerides for in situ storage rather than in phospholipids.

Sahoo et al., (2017) reported that the reduced glutathione (GSH) is the most important intracellular antioxidant [36]. Thanks to the activity of the glutathione peroxidase, GSH reacts with various oxidizing radicals (including hydrogen peroxide or reactive oxygen species ROS) to produce oxidized glutathione (GSSG) and reduced products. The system GSH-GSSG is one of the most powerful radical scavenger present in various cellular types and it is largely admitted that the increase of the ratio GSH/GSSG is a good indicator of the level of the oxidative status of the cell. According to Chen et al. (2013), the level of oxidized glutathione could be used as a marker of steatosis and oxidative stress [37]. In the present study, the ratio between LWL and HWL values for the oxidized glutathione is greatly lower than one indicating that high weight livers had higher levels of steatosis and oxidative stress than low weight livers. These results are in good accordance with previous observations focused on different proteic markers associated with the oxidative status of hepatocytes at the end of the overfeeding period [9].

Lactate is the product of the reduction of pyruvate, issued from glucose during glycolysis, in anaerobic conditions. In humans, Kalhan et al. (2011) showed that plasmatic lactate levels were higher when hepatic steatosis developed [38]. According to Murphy et al. (2001), hyperlactatemia could be the result of cellular hypoxia or hypoperfusion with resulting anaerobic metabolism [39]. Furthermore, McGarry et al. (2018) reported that hypoxia induces an increase in glycolytic activity through an increase in intermediate metabolites such as lactate [40]. Thus, the observed increase in lactate content in HWL at the end of the overfeeding period could reflect an orientation of the energetic metabolism of hepatocytes towards more anaerobism. This confirms previous reports indicating that high weight livers may suffer from hypoxia when steatosis develops [20].

The process of creatine synthesis is divided into two steps, catalyzed by the L-arginine:glycine amidinotransferase (AGAT) and the guanidinoacetate N-methyltransferase (GAMT) which takes place mainly in kidneys and liver, respectively [41]. Recently, Aljobaily et al. (2020) showed that a creatine supplementation in the feed of rats resulted in a decrease in the oxidative stress and damages in high weight livers [42]. This mainly reflects that creatine can play a direct role as an antioxidant [43]. According to Persky and Brazeau (2001), the benefits of creatine are generally attributed to the creatine-induced buffering of cellular ATP levels and its significant depletion will lead to accumulation of intracellular Ca^2 +^, formation of ROS and oxidative tissue damages [44]. In the present study, the level of creatine content was higher in HWL than in LWL. Probably because the oxidative stress occurred more in HWL than in LWL, HWL metabolism was more oriented towards a higher production of creatine to act as a supplementary antioxidant.

## Conclusion

The results of this study showed that the more or less important increase in the liver weight during an overfeeding period is associated with differences in the amounts of some hepatic metabolites. Livers with a relative low weight increase (final weight 550 g and 600 g) are characterized by a moderate utilization of the available quantity of glucose and therefore a lower lipid synthesis and more lipid export thanks to the higher quantity of choline available in the liver. This is associated with a possible decrease in the hepatic concentrations of triglycerides and cholesterol shown by a more marked presence of ethanolamine in those LWL. On the contrary, livers with a weight higher than 700 g are more characterized by a less efficient aerobic energetic metabolism (high amounts of lactate in HWL) associated with a higher level of oxidative stress reported by higher levels of glutathione oxidized and creatine.

Because it is known that different birds will differently support the metabolic shift imposed by the overfeeding, it is not totally surprising to finally see a more or less developed fatty liver. It is associated with a metabolomic profile illustrating a more or less active synthesis and exportation of lipids, or the beginning of their excessive accumulation associated with the beginning of cellular oxidation. Further studies will try to demonstrate if those metabolic states are transitory or definitive and therefore characteristics of the achievement of two different states of hepatic steatosis.

